# A comparative genomics framework for identifying historical population bottlenecks using olfactory receptor gene evolution

**DOI:** 10.64898/2026.08.03.742608

**Authors:** Kelly O’Regan, Louise Ryan, Graham Hughes

**Affiliations:** School of Biology and Environmental Science, University College Dublin, Ireland

**Keywords:** Conservation genomics, Population bottleneck, genetic diversity, mammals, chemosensory perception, olfactory receptors

## Abstract

Population bottlenecks reduce genetic diversity, increase the fixation of deleterious mutations and elevate extinction risk. Identifying lineages experiencing bottlenecks is a key goal of conservation genetics, facilitating the allocation of limited resources to at-risk species. Although whole-genome sequencing has improved bottleneck detection by reconstructing demographic history, these methods often require extensive population sampling, limiting their application. Previous studies of species showing population bottlenecks have reported an increased number of pseudogenes in the olfactory receptor (OR) gene family, however whether such evolutionary dynamics can be used as comparative biomarkers of genomic decline remains unknown. By quantifying the number of lineage-specific duplication and pseudogenization events, we introduce the duplication-to-loss ratio (DLR), a comparative metric exploring the rate at which chemosensory gene loss is offset by the generation of novel receptors. We characterize the chemosensory repertoires of 21 felid species, including species with known historical bottlenecks, to establish the utility of this DLR metric. Subsequently, we evaluate its usage across additional mammalian families, specifically Ursidae and Pinnipedia, to determine its utility beyond Felidae. Our DLR metric recovers several felid species with a history of genomic decline, including cheetah (*Acinonyx jubatus*) and black-footed cat (*Felis nigripes*), as well as the giant panda (*Ailuropoda melanoleuca*), polar bear (*Ursus maritimus*), Hawaiian monk seal (*Neomonachus schauinslandi*) and northern elephant seal (*Mirounga angustirostris*). Our results demonstrate the utility of the OR gene repertoire as a scalable, robust biomarker for identifying comparative population decline, prioritising species for conservation genomic investigation using only the reference genome.

## Introduction

Population bottlenecks, where a sharp reduction in population size results in reduced genetic diversity and increased extinction risk, have been observed in a wide range of species (Nei *et al*., 1975; Frankham, 2003; van Oosterhout *et al*., 2026). A low effective population size (*N_e_*) accelerates the impact of genetic drift, leading to a rapid loss of genetic variation and facilitating the fixation of deleterious alleles through inbreeding and relaxed selection. Historical shifts in climate, population fragmentation and intense anthropogenic activity have all led to demographic decline (Hewitt, 2000; Ceballos *et* al., 2015; Haddad *et al*., 2015). Identifying lineages experiencing such a decline is a key goal of conservation genetics, as it allows the allocation of limited resources to the most at-risk species (Allendorf *et al*., 2010; Garner *et al* 2015; Hogg, 2023). While whole-genome data is increasingly used to reconstruct demographic history, limitations including low population density or sample size, sample collection logistics, and historical selective sweeps can impact efforts in conservation genetics, highlighting the need for complementary genomic approaches capable of identifying historical decline from limited resources (Luikart *et al*., 1998; Peery *et al.,* 2012; Kardos *et* al., 2021; Clark *et* al., 2024; Fedorca *et al*., 2024; Pegueroles *et al*., 2024).

Chemosensory multigene families are some of the largest and most rapidly evolving gene families in mammals, consisting of olfactory receptors (ORs) and trace amine associated receptors (TAARs) for odorant binding, vomeronasal receptors (V1R/V2Rs) for pheromone detection and taste receptors (TAS1R/TAS2Rs) for gustatory perception (Nelson *et al*., 2001; Sullivan, 2002; Mombaerts, 2004; Yang *et al*., 2005; Niimura and Nei, 2006; Liberles, 2009; Meyerhof and Korsching, 2009; Behrens and Meyerhof). Chemosensory genes are dispersed across multiple chromosomes and are, with the exception of T1Rs and V2Rs, generally encoded by a single exon (Niimura and Nei, 2003; Malnic *et al*., 2004). Chemosensory genes can also differ in frequency considerably across lineages due to ecological niche adaptation and genome evolution. These families evolve through a process of gene birth-and-death, where duplications result in novel receptors and loss-of-function mutations result in the accumulation of pseudogenes (Nei and Rooney, 2005; Dong *et al*., 2009; Hughes *et al*., 2018). Such duplicate genes are often under reduced selective constraint, making them especially susceptible to the fixation of loss-of-function mutations following a reduction in *N_e_*.

Reductions in genetic diversity due to historical population bottlenecks have been observed in several species including *Acinonyx jubatus* (cheetah; O’Brien *et al*., 1983; Dobrynin *et al*., 2015), *Mirounga angustirostris* (elephant fur seal; Hoelzel *et al*., 2024), *Enhydra lutris* (sea otter; Beichman *et al*., 2019) and *Moschus anhuiensis* (musk deer; Chen *et al*., 2025). A common observation made in several of these species is an increase in the number of non-functional OR genes. This observation has also been recorded in certain extinct species (specifically Wrangel island woolly mammoth (*Mammuthus primigenius*; Rogers and Slatkin, 2017) and thylacine (*Thylacinus cynocephalus*; Salve and Vijay, 2025). These studies suggest that OR pseudogenization is a recurring consequence of demographic decline, capturing evidence of historical events such as genetic bottlenecks. However, as loss of OR function has generally been reported as a secondary observation rather than the primary focus of these studies, it remains unclear whether they can be used as a potential comparative genomics biomarker. As the total number of OR pseudogenes can be confounded by historical losses in ancestral lineages (Hughes *et al*., 2018), pseudogene counts alone cannot distinguish between inherited or unique loss events across species.

Here, we investigate whether lineage-specific chemosensory gene evolution, specifically OR genes, provides a useful biomarker of historical genomic decline. By comparing the number of OR duplications generating new functional receptors with pseudogenizations unique to individual lineages, we introduce the duplication-to-loss ratio (DLR) to determine the rate at which gene loss is offset by the birth of new receptors. We first characterize the chemosensory repertoires of species within the Felidae family to determine whether DLR identifies species with documented histories of reduced genetic diversity. By applying this framework to species within Ursidae and Pinnipedia, we evaluate the broader use of DLR in different mammalian lineages. Our comparative analysis shows that the OR repertoire and its evolutionary dynamics serves as a useful metric for revealing at-risk species requiring further conservation genomics investigation.

## Materials and Methods

### Felid genome assemblies and completeness

A total of 22 felid genomes, including cheetah, were downloaded from RefSeq (Goldfarb et al, 2025; **Table 1**). Pre-determined completeness scores for 20 genomes were collated from previous studies (**Table 1**). Genomes with BUSCO (Seppey *et al*., 2019) completeness scores below 85% were excluded to minimise biases associated with incomplete genome assemblies and avoid skewed inference of sensory gene repertoire composition. This range of species spanned the Felidae tree and was used to establish the background felid sensory repertoire for comparative analysis.

**Table 1.**
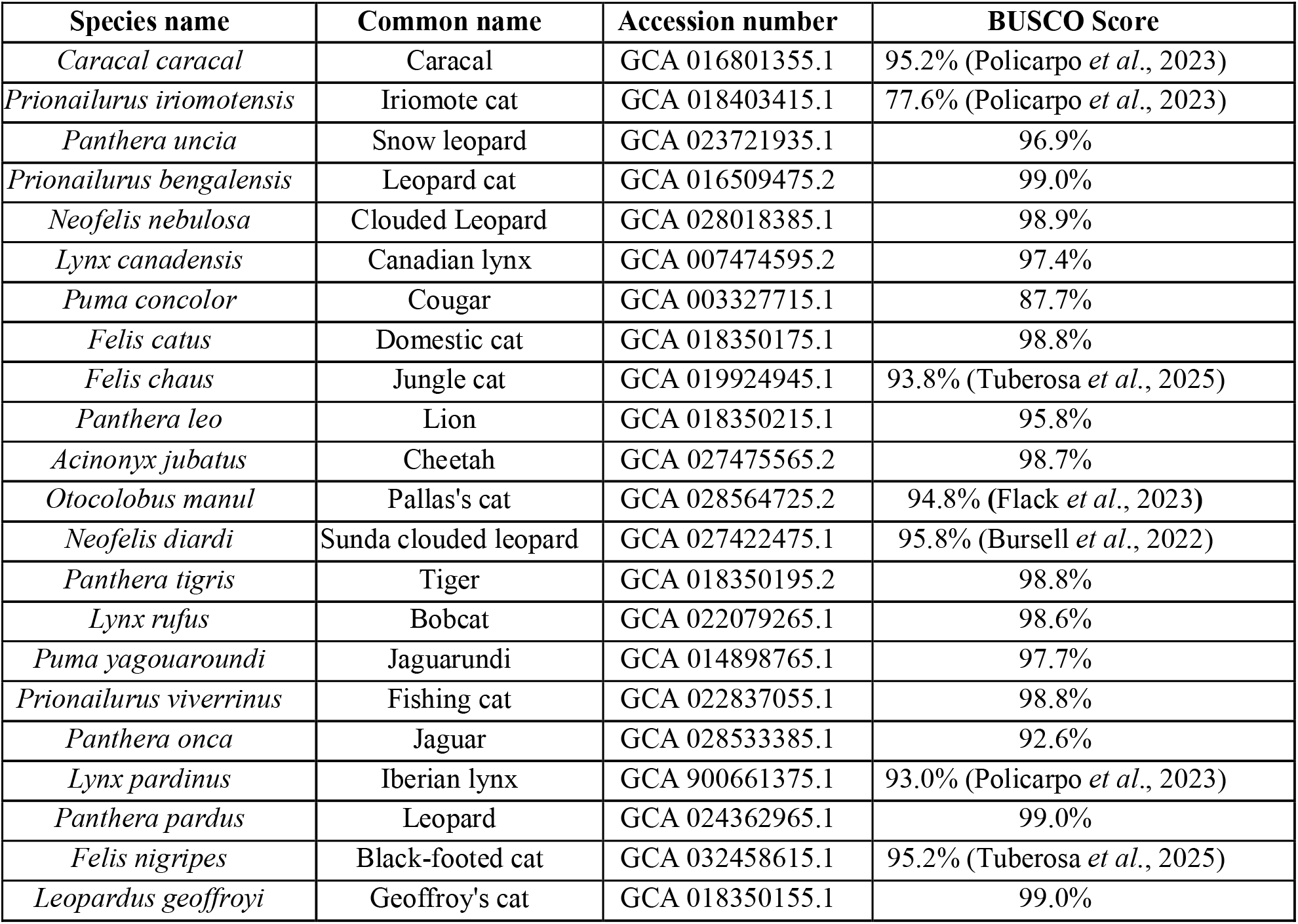
Target Felid species. The sensory repertoire from a total of 22 felid species were explored. Accession numbers and BUSCO genome completeness scores are shown.

| Species name | Common name | Accession number | BUSCO Score |
| --- | --- | --- | --- |
| *Caracal caracal* | Caracal | GCA 016801355.1 | 95.2% (Policarpo *et al.*, 2023) |
| *Prionailurus iriomotensis* | Iriomote cat | GCA 018403415.1 | 77.6% (Policarpo *et al.*, 2023) |
| *Panthera uncia* | Snow leopard | GCA 023721935.1 | 96.9% |
| *Prionailurus bengalensis* | Leopard cat | GCA 016509475.2 | 99.0% |
| *Neofelis nebulosa* | Clouded Leopard | GCA 028018385.1 | 98.9% |
| *Lynx canadensis* | Canadian lynx | GCA 007474595.2 | 97.4% |
| *Puma concolor* | Cougar | GCA 003327715.1 | 87.7% |
| *Felis catus* | Domestic cat | GCA 018350175.1 | 98.8% |
| *Felis chaus* | Jungle cat | GCA 019924945.1 | 93.8% (Tuberosa *et al.*, 2025) |
| *Panthera leo* | Lion | GCA 018350215.1 | 95.8% |
| *Acinonyx jubatus* | Cheetah | GCA 027475565.2 | 98.7% |
| *Otocolobus manul* | Pallas's cat | GCA 028564725.2 | 94.8% (Flack *et al.*, 2023) |
| *Neofelis diardi* | Sunda clouded leopard | GCA 027422475.1 | 95.8% (Bursell *et al.*, 2022) |
| *Panthera tigris* | Tiger | GCA 018350195.2 | 98.8% |
| *Lynx rufus* | Bobcat | GCA 022079265.1 | 98.6% |
| *Puma yagouaroundi* | Jaguarundi | GCA 014898765.1 | 97.7% |
| *Prionailurus viverrinus* | Fishing cat | GCA 022837055.1 | 98.8% |
| *Panthera onca* | Jaguar | GCA 028533385.1 | 92.6% |
| *Lynx pardinus* | Iberian lynx | GCA 900661375.1 | 93.0% (Policarpo *et al.*, 2023) |
| *Panthera pardus* | Leopard | GCA 024362965.1 | 99.0% |
| *Felis nigripes* | Black-footed cat | GCA 032458615.1 | 95.2% (Tuberosa *et al.*, 2025) |
| *Leopardus geoffroyi* | Geoffroy's cat | GCA 018350155.1 | 99.0% |

### Chemosensory gene annotation

The chemosensory gene repertoire of each species, specifically olfactory receptors (OR), vomeronasal receptors (V1R/V2R), taste receptors (T1R/T2R) and trace amine-associated receptors (TAAR), were mined and annotated using the *Sensommatic* pipeline (Ryan *et al*., 2024). *Sensommatic* utilizes BLAST (Altschul *et al*., 1990) and Augustus (Stanke *et al*., 2004) to identify sensory genes in a reference genome using a set of reference mammal sequences. Genes were classified using default parameters, with a lack of in-frame stop codons and minimum length of 860bp in single exon receptor genes (OR, TAAR, V1R, TAS1R), 1000bp for TAS1Rs, and 2000bp for V2Rs used as thresholds for functionality. To ensure no receptors were missed, all annotated sequences were combined and independently mapped to each genome using BLAST, with putative receptor sequences not already contained in the *Sensommatic* outputs subsequently extracted.

### Comparison of sensory genes across species

Raw counts of functional and non-functional genes, as well as the ratio of functional and non-functional receptors per repertoire were used to compare across species. Principal components analysis on the proportion of total functional genes per gene family, as well as subfamilies within the OR gene repertoire, were used to explore how similar/dissimilar each felid species was to the others. The impact of each species on the overall clustering pattern was observed using a ‘leave-one-out’ approach. For this, the total variance accounted for by PC1 and PC2 was initially computed for the proportion of functional genes across species. Individual species were then sequentially removed from the count data, with variance subsequently re-estimated and compared to identify possible outliers, with large variation suggesting that specific species heavily impacts the patterns observed.

### Phylogenetic inference of sensory gene trees

For each gene subfamily (OR: OR1/3/7, OR2/13, OR4, OR5/8/9, OR6, OR10, OR11, OR12, OR14, OR51, OR52, OR55, OR56, OR undefined (OR genes not placed in a specific family); V1R: VN1R1-5, VN1R43, VN1R90, VN1R94; V2R:V2R1, V2R24, V2R26, V2R35, V2R116, V2R309; TAS1R: TAS1R1-3, TAS2R: TAS2R1-60, TAAR: TAAR1-9), multiple sequence alignments of functional and non-functional receptor nucleotide sequences were generated using MACSE (Ranwez *et al*., 2018). Gene trees for each subfamily were inferred using IQTREE2 (Minh *et al*., 2020) with the best-fit model of sequence evolution (Kalyaanamoorthy *et al*., 2017). Given their large size, OR gene families 2/13 and 5/8/9 were aligned as separate families (2, 13, 5, 8, 9). Global patterns of Felidae sensory gene evolution were established against a composite phylogenetic species tree (Li *et* al., 2016; Foley *et al*., 2023). Branch lengths for this species tree were estimated using the Grafen method implemented in the *ape R* package (Grafen 1989; Paradis *et al*., 2004).

### Lineage-specific gene duplication-and-loss

Lineage-specific duplications for each gene family per lineage were identified via gene tree-species tree reconciliation as implemented in NOTUNG (Chen *et al*., 2000; Stolzer *et al*. 2012). This was done using the inferred gene trees and the composite species tree phylogeny. Only instances of duplication where both paralogs maintained functionality were considered (**Figure 1a**). Lineage-specific losses, determined as instances where a specific gene in a target species has become pseudogenized relative to orthologs in its sister taxa, were also identified for each species using Orthosnap (Steenwyk *et al*., 2022; **Figure 1b**). Specifically, all single-copy orthologous monophyletic subgroups per gene tree were identified, with a loss considered lineage-specific if it occurred in only one species within that subgroup.

**Figure 1.**
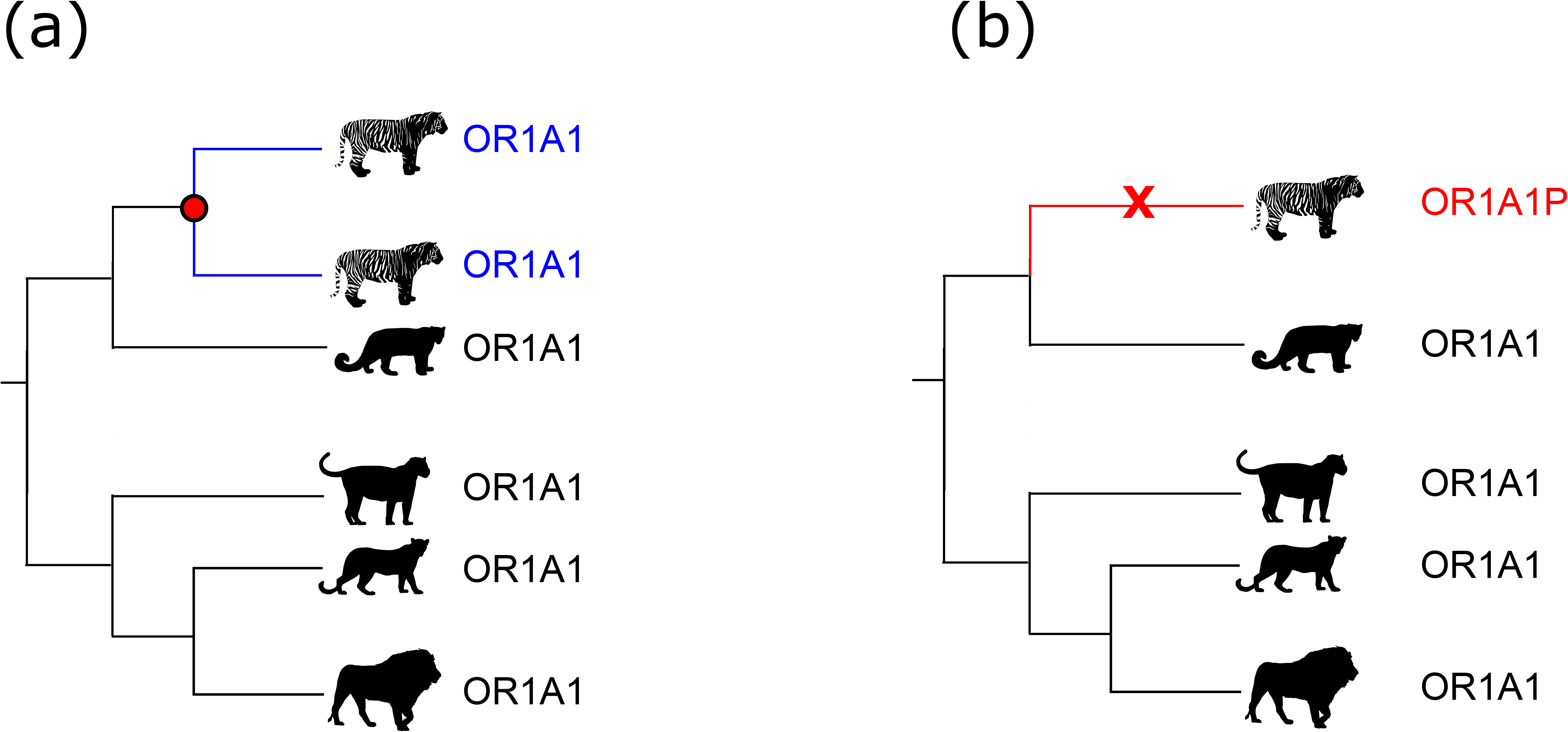
Lineage-specific chemosensory evolution. Chemosensory genes that evolve under the birth-and-death model show lineage-specific duplications and losses. Duplications within a species resulting in two functional paralogs (a) and unique pseudogenization events (b) were the focus of this study.

As a higher rate of pseudogenization was our metric for uncovering instances of historical loss in genetic diversity, we calculated the ratio of lineage-specific duplications, *D,* to lineage-specific losses, *L* (D/L), using this duplication-to-loss ratio (DLR) to determine if sensory gene loss was occurring at a higher frequency than birth. This metric requires at least one lineage-specific pseudogenization per species and was applied to both the combined repertoire of all sensory genes and OR genes only. A DLR value greater than one suggests a high turnover of new functional receptors, exceeding that of loss-through-pseudogenization events, a DLR less than one implies gene loss is not being offset by gene duplications, and a value of one suggests an evolutionary balance within the chemosensory gene family. Our DLR metric can be extended to both account for instances of no duplications and be centred around zero using log(*D*+1)-log(*L*+1); (log-DLR), where log-DLR>0 indicates a higher rate of duplications than losses, log-DLR<0 a higher rate of loss compared to duplications, and log-DLR=0 indicating equal rates of evolution.

### Application to additional species

To further explore the use of chemosensory genetic biomarkers, the DLR ratio was applied to additional orders with known instances of historical reductions in effective population sizes. Specifically, six species within the Ursidae family: giant panda (*Ailuropoda melanoleuca*), spectacled bear (*Tremarctos ornatus*), American black bear (*Ursus americanus*), brown bear (*Ursus arctos*), polar bear (*Ursus maritimus*), Asian black bear (*Ursus thibetanus*; **Supplementary Table 1**) and 14 species in Pinnipedia: Antarctic fur seal (*Arctocephalus gazella*), Guadalupe fur seal (*Arctocephalus Townsendi*), northern fur seal (*Callorhinus ursinus*), steller sea lion (*Eumetopias jubatus*), grey seal (*Halichoerus grypus*), Weddel seal (*Leptonychotes weddellii*), northern elephant seal (*Mirounga angustirostris*), southern elephant seal (*Mirounga leonina*), Hawaiian monk seal (*Neomonachus schauinslandi*), Pacific walrus (*Odobenus rosmarus divergens*), harbour seal (*Phoca vitulina*), Saimaa ringed seal (*Pusa hispida saimensis*), Baikal seal (*Pusa sibirica*), Californian sea lion (*Zalophus californianus*) were explored. Ursidae was chosen as both the giant panda and polar bear have undergone several extreme population bottlenecks (Zhang *et al*., 2007; Zhu *et al*., 2013; Liu *et* al, 2015; Maduna *et al*., 2021; Lan *et al*., 2022), while several pinnipeds have undergone known population bottlenecks (Stoffel *et al*., 2018). The DLR and log-DLR values in OR and total sensory genes were determined using our methods described above.

## Results

### Felid sensory gene repertoires

Only one genome out of 22, Iriomote cat (*Prionailurus iriomotensis*), had a BUSCO completeness score below 85% excluding it from downstream analysis (**Table 1**). A total of 25,199 genes, consisting of 23,048 ORs, 1136 V1Rs, 207 V2Rs, 68 TAS1Rs, 487 TAS2Rs, 253 TAARs were identified across 21 species (mean ORs: 1098; V1Rs: 54; V2Rs: 10; TAS1Rs: 3; TAS2Rs: 23; TAARs: 12), with a mean of 71.7% of sensory receptors showing functionality (**Table 2**). The highest and lowest number of functional sensory genes were found in leopard (*Panthera pardus*; n=1004) and cougar (*Puma concolor*; n=664) respectively, while snow leopard (*Panthera uncia*) and Geoffroy’s cat (*Leopardus geoffroyi*) had the highest (n=757) and lowest (n=250) pseudogene counts, reflecting differences in ecological adaptations and sensory niches.

**Table 2.** Total number of sensory genes. The number of sensory genes, functional and non-functional, as well as their mean values across 21 felid species are displayed.

| Gene family | Total | Mean | Functional | Mean functional | Pseudogenes | Mean pseudogenes |
| --- | --- | --- | --- | --- | --- | --- |
| OR | 23,048 | 1,098 | 16,898 | 805 | 6,150 | 293 |
| V1R | 1,136 | 54 | 548 | 26 | 588 | 28 |
| V2R | 207 | 10 | 95 | 10 | 112 | 5 |
| TAS1R | 68 | 3 | 45 | 2 | 23 | 1 |
| TAS2R | 487 | 23 | 297 | 14 | 190 | 9 |
| TAAR | 253 | 12 | 195 | 9 | 58 | 3 |
| Total | 25,199 | 1,200 | 18,078 | 861 | 7,121 | 339 |

### Sensory repertoire composition across felidae

The proportion of functional genes per receptor family and subfamily was used to measure the retention of functionality for that sensory modality per species. Jaguar (*Panthera onca*) showed the lowest proportion of functional genes for the OR (50%), TAAR (50%) and V1R (35%) gene families, while Iberian lynx (*Lynx pardinus*, 40%) and cougar (45%) had the lowest proportion of functionality for TAS1R and TAS2R families, respectively. Within the OR gene family, jaguar also had the lowest proportion for eight of the 13 gene subfamilies (**Supplementary Table 2**), followed by black-footed cat (Felis nigripes; four subfamilies) and Iberian lynx (one subfamily). Principal component analyses both across all sensory families (PC1+PC2 accounting for 72.1% of total variance) and within ORs (PC1+PC2 accounting for 87.8% of total variance) largely reflected this disparity in functionality across species (**Supplementary Figure 1-2**). When using the ‘leave-one-out’ approach, the removal of *P. onca* had the largest impact on PC1 while caracal (*Caracal caracal*) had the largest impact on PC2, suggesting highly divergent chemosensory gene repertoire profiles within these species (**Supplemental Table 3**).

### Lineage-specific gain and loss across felid species

Gene tree-species tree reconciliation highlighted a mean of 44 duplications per species, with caracal and cougar having the highest (n=119) and lowest (n=9) frequency of gene births through duplication. Only 21 duplications where both paralogs are functional were found across all species, with the fewest found in the black-footed cat. The mean number of lineage-specific gene losses was 13 per species, with jaguar (n=53) showing the highest frequencies and both jungle cat (*Felis chaus*) and bobcat (*Lynx rufus*; n=4) showing the lowest. Using a ratio threshold of 1, the ratio of lineage-specific gene duplications to gene losses (DLR) in both the OR gene and full chemosensory gene repertoire highlighted eight species (Sunda clouded leopard (*Neofelis diardi*), Canadian lynx (*Lynx canadensis*), Pallas’ cat (*Otocolobus manul*), jaguar, black-footed cat, cougar, cheetah and snow leopard) as losing more sensory/OR genes through pseudogenization than they are gaining through duplication (DLR<1; **Table 3**, **Figure 2, Supplementary Table 2**), reflecting documented histories of a loss in genetic diversity. These results were also reflected in the log-DLR values across species (**Supplemental Figure 3)**.

**Figure 2.**
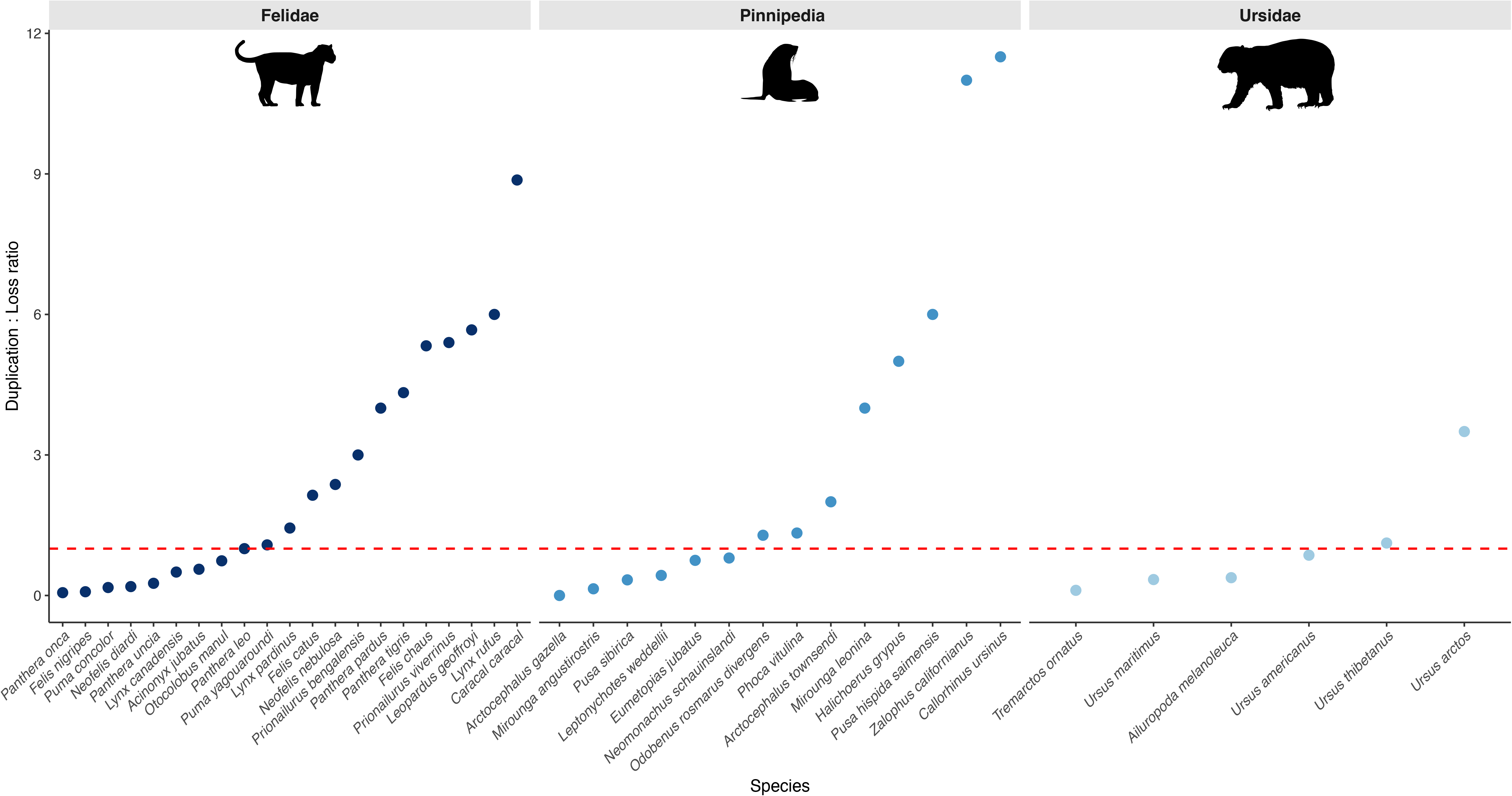
Duplication-to-loss (DLR) ratios. The ratio of OR gene duplication to gene loss was used as a metric to determine species with historical decreases in genetic diversity in three mammal groups. A threshold of 1 (red dashed line) was applied, with lineages below 1 representing at-risk species.

**Table 3.** Ratio of OR gene duplications to loss. The ratio of functional gene duplications, where both paralogs have retained function, to lineage-specific gene loss events (DLR) were used to determine if functional OR genes are reducing in different species.

| Species | Common name | OR Duplications | OR Loss | DLR | log-DLR |
| --- | --- | --- | --- | --- | --- |
| **Felidae** |  |  |  |  |  |
| *Panthera onca* | Jaguar | 3 | 53 | 0.06 | -1.13 |
| *Felis nigripes* | Black-footed cat | 2 | 24 | 0.08 | -0.92 |
| *Puma concolor* | Cougar | 4 | 23 | 0.17 | -0.68 |
| *Neofelis diardi* | Sunda clouded leopard | 3 | 16 | 0.19 | -0.63 |
| *Panthera uncia* | Snow leopard | 6 | 23 | 0.26 | -0.54 |
| *Lynx canadensis* | Canadian lynx | 4 | 8 | 0.5 | -0.26 |
| *Acinonyx jubatus* | Cheetah | 9 | 16 | 0.56 | -0.23 |
| *Otocolobus manul* | Pallas's cat | 17 | 23 | 0.74 | -0.12 |
| *Panthera leo* | Lion | 9 | 9 | 1 | 0 |
| *Puma yagouaroundi* | Jaguarundi | 14 | 13 | 1.08 | -1.13 |
| *Lynx pardinus* | Iberian lynx | 13 | 9 | 1.44 | 0.15 |
| *Felis catus* | Domestic cat | 15 | 7 | 2.14 | 0.3 |
| *Neofelis nebulosa* | Clouded Leopard | 19 | 8 | 2.37 | 0.35 |
| *Prionailurus bengalensis* | Leopard cat | 12 | 4 | 3 | 0.41 |
| *Panthera pardus* | Leopard | 28 | 7 | 4 | 0.56 |
| *Panthera tigris* | Tiger | 26 | 6 | 4.33 | 0.59 |
| *Felis chaus* | Jungle cat | 16 | 3 | 5.33 | 0.63 |
| *Prionailurus viverrinus* | Fishing cat | 27 | 5 | 5.4 | 0.67 |
| *Leopardus geoffroyi* | Geoffroy's cat | 34 | 6 | 5.67 | 0.7 |
| *Lynx rufus* | Bobcat | 18 | 3 | 6 | 0.68 |
| *Caracal caracal* | Caracal | 71 | 8 | 8.87 | 0.9 |
| **Ursidae** |  |  |  |  |  |
| *Tremarctos ornatus* | Spectacled bear | 14 | 131 | 0.11 | -0.94 |
| *Ursus maritimus* | Polar bear | 10 | 29 | 0.34 | -0.44 |
| *Ailuropoda melanoleuca* | Giant Panda | 45 | 118 | 0.38 | -0.41 |
| *Ursus americanus* | American Black bear | 18 | 21 | 0.86 | -0.06 |
| *Ursus thibetanus* | Asian black bear | 84 | 75 | 1.12 | 0.05 |
| *Ursus arctos* | Brown bear | 28 | 8 | 3.5 | 0.51 |
| **Pinnipedia** |  |  |  |  |  |
| *Arctocephalus gazella* | Antarctic fur seal | 0 | 25 | 0 | -1.41 |
| *Mirounga angustirostris* | Northern elephant seal | 1 | 7 | 0.14 | -0.6 |
| *Pusa sibirica* | Baikal seal | 1 | 3 | 0.33 | -0.3 |
| *Leptonychotes weddellii* | Weddell seal | 3 | 7 | 0.43 | -0.3 |
| *Eumetopias jubatus* | Steller sea lion | 3 | 4 | 0.75 | -0.1 |
| *Neomonachus schauinslandi* | Hawaiian monk seal | 8 | 10 | 0.8 | -0.09 |
| *Odobenus rosmarus divergens* | Pacific walrus | 9 | 7 | 1.29 | 0.1 |
| *Phoca vitulina* | Harbor seal | 4 | 3 | 1.33 | 0.1 |
| *Arctocephalus townsendi* | Guadalupe fur seal | 4 | 2 | 2 | 0.22 |
| *Mirounga leonina* | Southern elephant seal | 4 | 1 | 4 | 0.4 |
| *Halichoerus grypus* | Grey seal | 5 | 1 | 5 | 0.48 |
| *Pusa hispida saimensis* | Saimaa ringed seal | 6 | 1 | 6 | 0.54 |
| *Zalophus californianus* | California sea lion | 11 | 1 | 11 | 0.78 |
| *Callorhinus ursinus* | Northern fur seal | 23 | 2 | 11.5 | 0.9 |

### Application to Ursidae and Pinnipedia

A total of 11,186 sensory genes (ORs: 10,640, TAARs: 66, TAS1Rs: 19, TAS2Rs: 132, V1Rs: 286, V2Rs: 24) and 11,816 sensory genes (ORs: 10,878, TAARs: 110, TAS1Rs: 29, TAS2Rs: 304, V1Rs: 433, V2Rs: 62) were recovered across Ursidae and Pinnipedia, respectively (**Supplementary Table 4**). Within both orders, the highest and lowest number of functional receptors were found in the brown bear and giant panda within Ursidae, and Saimaa ringed seal and Antarctic fur seal in Pinnipedia. The mean number of lineage-specific duplications in Ursidae was 214 per species, which is higher than the rates observed in both Felidae and Pinnipedia (mean of 48 per species) (**Supplementary Table 5**). When comparing duplication rate to loss, four ursid species and six pinnipeds had a DLR<1 (**Table 3, Figure 2, Supplemental Figure 3**).

## Discussion

The loss of genetic diversity such as that which follows a population bottleneck substantially increases extinction risk through a reduction in variation and increase of deleterious mutations. Identifying signatures of historical loss diversity represents an important step in conservation genomics, as it can facilitate the prioritization of at-risk species for preservation initiatives and programs. Here, we have shown that lineage-specific chemosensory gene loss, specifically within the olfactory receptor (OR) gene repertoire, can be used to identify species with a history of genetic decline. Using three mammalian orders, we have demonstrated that the ratio of OR duplications to losses consistently recovers species with known reductions in genetic diversity relative to their sister taxa. As the generation of reference-quality genomes in conservation genomics becomes increasingly more common, this approach offers a rapid screening tool to identify lineages that warrant further exploration.

### Chemosensory evolution as a useful biomarker for identifying historical loss of diversity

Chemosensory receptors, including olfactory receptors (ORs), evolve through gene birth-and-death, with duplications leading to new paralogous receptors. Following duplication, one paralog may experience relaxed functional constraint leading to an accumulation of loss-of-function mutations (pseudogenes). In populations with a large effective population size (*N*_e_), these pseudogenes may be specific to an individual, never reaching fixation and staying at low frequencies. However, a severe or prolonged reduction in *N*_e_ may lead to the rapid fixation of these loss-of-function mutations. Consequently, large multigene families, such as ORs, are expected to be particularly susceptible to pseudogenizations than single-copy genes during a bottleneck. This hypothesis is supported by observations of increased pseudogenization in OR genes in several case studies (Rogers and Slatkin, 2017; Beichman *et al*., 2019; Chen *et al*., 2025; Salve and Vijay, 2025). By comparing the frequency of lineage-specific duplications with lineage-specific pseudogenization, we determined the rate at which chemosensory gene loss is being offset by the generation of new functional genes. Our results suggest that the OR repertoire alone has enough signal associated with historical decline in genetic diversity without incorporating additional chemosensory gene families.

One caveat here is that ecological adaptation itself is a major driver of chemosensory gene evolution. The expansion and contraction of these gene families can occur alongside changes in life-history traits including diet, habitat, rhythmic activity and sociality (Hughes *et al*., 2018). An increase in the number of pseudogenes may represent modifications in sensory ecology rather than population dynamics alone. While calculating lineage-specific gain and loss of function in a phylogenetic and comparative genomics framework can mitigate this, it is unlikely to fully disentangle sensory adaptation with mammalian demography. We therefore consider the DLR as an additional indicator of historical loss of genetic diversity rather than a direct measure of past population bottlenecks.

### Application of duplication-to-loss ratio to Felidae

Using the DLR as our key metric, we identified several species where the rate of lineage-specific OR pseudogenization exceeded the generation of new functional receptors through duplication (DLR<1). Several felid species with documented or suspected histories of demographic decline or reductions in genetic diversity were highlighted including cheetah (O’Brien *et al*., 1987; O’Brien *et al*., 2017), black-footed cat (Grant *et al*., 2026), cougar (Gustafson *et al*., 2022; Mac Allister *et al*., 2024), Sunda clouded leopard (Bursell *et al.,* 2022) and Pallas’ cat (Bubenikova *et al*., 2024). The snow leopard, which also fell below our threshold, has experienced continuously small population sizes and low *N*_e_ values throughout its evolutionary history (Solari *et al*., 2025), hinting at a possible confounding factor that may influence OR repertoire evolution and the fixation of OR pseudogenes. While species with DLR>1 have experienced some habitat fragmentation or local population decline (e.g. caracal; Kyriazis *et al*., 2024), others are considered stable or increasing by the International Union for Conservation of Nature (IUCN)(e.g. Iberian lynx, bobcat (Miller-Butterworth *et al*., 2021), Leopard cat (*P. bengalensis*) suggesting high DLRs are consistent with a lack of long-term erosion of genetic diversity (https://www.iucnredlist.org/search?taxonomies=101738&searchType=species; **Supplemental Figure 3**). The most unexpected result was the exceptionally low DLR observed in the jaguar. Our analyses of jaguar highlighted how much it differed to other species based on the proportion of functional receptors and due to it having the greatest rate of OR loss, despite genome-wide studies generally highlighting their moderate-to-high levels of variability (Valdez *et al*., 2015; Wultsch *et al*., 2016; Lorenzana *et al*., 2020). Despite this, whole-genome studies show lower rates of diversity as well as instances of large reductions in *N*_e_ over the past 1-2 million years during the Pleistocene (Lorenzana *et al*., 2022) and low levels of heterozygosity in some distinct *P. onca* populations (Meißner *et al*., 2025). This explains certain aspects of our results; however, it is also possible that differing evolutionary pressures and ecological specialization (Figueiró *et al*., 2017) have driven the increased rate of OR gene loss. Additionally, while genome quality and sample source may also influence our analysis (e.g. jaguar genome comes from a captive individual in Heuston zoo, BioSample: SAMN14122073), the relatively high BUSCO scores and broad signal of gene loss across other chemosensory gene families suggests this impact is minimal. Nonetheless, the recovery of multiple felid species with independent genomic evidence of historical losses in genetic diversity, irrespective of convergent or divergent evolutionary trajectories, suggests that increased rates of OR genes pseudogenization reflects a wider reduction in *N*_e_ making our metric a useful tool for future conservation studies.

### Exploring OR gene loss in Ursidae and Pinnipedia

To determine whether an increase in OR gene loss reflected historical losses of diversity outside of Felidae, we applied our method to Ursidae and Pinnipedia, two additional mammalian families with well-known demographic histories. Within Ursidae, four of the six species showed a DLR below one. This included polar bear and giant panda, both of which show historical reductions in *N*_e_ and reduced genetic diversity (Zhu *et al*., 2013; Liu *et al*., 2015). The spectacled bear had the lowest DLR, consistent with previous studies highlighting low genetic diversity, small *N*_e_, and population fragmentation (Ruiz-Garcia, 2003; Ruiz-Garcia *et al*., 2005; Cueva *et al*., 2018), while the brown bear had the highest ratio consistent with its widespread geographic distribution and relatively large *N*_e_ (Miller *et al*., 2012; Kumar *et al*., 2017; Armstrong *et al*., 2022).

Our analysis of pinnipeds similarly recovered species with established histories of genetic decline. As with the cheetah in Felidae, the northern elephant seal is a common example of an extreme population bottleneck, reflected here in its severely low DLR (Weber *et* al., 2000; Abadía-Cardoso *et al*., 2017; Hoelzel *et al*., 2024; Hoffman *et al*., 2024). Reports of historical bottlenecks were also reflected in the DLR values of Hawaiian monk seal, steller sea lion, Antarctic fur seal and Baikal seal (Hoffman *et* al., 2006; Humble *et al*., 2018; Stoffel *et al*., 2018; Yakupova *et al*., 2023). A complete lack of new functional OR genes through duplication in the Antarctic fur seal was observed, suggesting a relaxed evolutionary pressure on the sense of olfaction relative to its sister taxon, Guadalupe fur seal. Both the northern fur seal and Californian sea lion had the highest DLR. While commercial hunting in recent centuries has severely reduced population sizes in both species (Zavala-Gonzalez and Mellink, 2000), the northern fur seal has retained multiple breeding populations and gene flow (Dickerson *et al*. 2010; Stoffel *et al*., 2018) while the Californian sea lion has shown rapid population recovery (Laake *et al*., 2018, Stoffel *et al*., 2018), suggesting the DLR metric is not impacted by recent population crashes.

## Conclusion

Historical losses of genetic diversity leave lasting genomic signatures, the discovery of which is a key component in conservation genomics. Here we have developed a simple comparative genomics framework for identifying species with historical reductions in effective population sizes. Using three different mammalian groups, with divergent life-history and evolutionary trajectories, we show the utility of the chemosensory genes, specifically OR genes, as a biomarker for population health. As the number of available reference-quality genomes rapidly increases, we believe this approach can be used to rapidly screen for highlighting potential at-risk species. While we looked at a specific mammalian order, future work exploring multiple genomes and estimates of population diversity will expand on the utility of this method across vertebrates.

## Acknowledgments

We thank Dr. Megan Power and Dr. John Finarelli for their suggestions and input in this project. All computational analyses were carried out using the UCD Sonic HPC.

## Author Contributions

The study was conceived by GMH. Methodology and study design, K.O’R., L.R., G.M.H. Sensory receptor gene mining and classification, L.R. Data analysis, interpretation, and visualisation, K.O’R., L.R., G.M.H. Writing, reviewing, and editing the manuscript, K.O’R. and G.M.H. Supervision and project oversight, G.M.H. All authors have read and approved the final manuscript.

## Supplementary material

Supplementary material is available at XXXXXX.

## Data availability

All datasets will be made available online through DRYAD at: [DOI Listed here if accepted]

## Supplementary Figures

**Supplementary Figure 1. Principal component analysis (PCA) of functional chemosensory genes.** A PCA analysis of OR, V1R, V2R, TAAR, TAS1R and TAS2R receptors (72.15 variance) highlights diversity in functional chemoreceptor genes.

**Supplementary Figure 2. Principal component analysis (PCA) of functional OR genes.** PCA of the functional OR gene repertoire in Felidae (88% variance) reveals several outlier species.

**Supplementary Figure 3. Olfactory receptor Log-DLR.** The duplication-to-loss (DLR) ratio and Log-DLR was applied to species within Felidae, Pinnipedia and Ursidae to highlight potential at-risk species. The IUCN status of each species is also displayed.

## Supplementary Tables

**Supplementary Table 1. Ursidae and Pinnipedia list of species**. All species from Ursidae and Pinnipedia, including their genome accession numbers, are displayed.

**Supplementary Table 2. Chemosensory repertoire sizes in Felidae.** The number of chemosensory receptors, functional and non-functional, as well as lineage-specific duplications and losses per species are displayed.

**Supplementary Table 3. PCA variation.** PCA analysis of Felidae using the leave-one-out approach highlight the species with the highest impact.

**Supplementary Table 4. Chemosensory repertoire sizes in Ursidae and Pinnipedia.** The number of chemosensory receptors, functional and non-functional, for each species within both Ursidae and Pinnipedia are displayed.

**Supplementary Table 5. Duplications and losses within Ursidae and Pinnipedia**. The number of lineage-specific duplication and loss events per species are displayed.

