## Supplementary figures and images for "A comparative genomics framework for identifying historical population bottlenecks using olfactory receptor gene evolution"

### Supplementary Figure 1

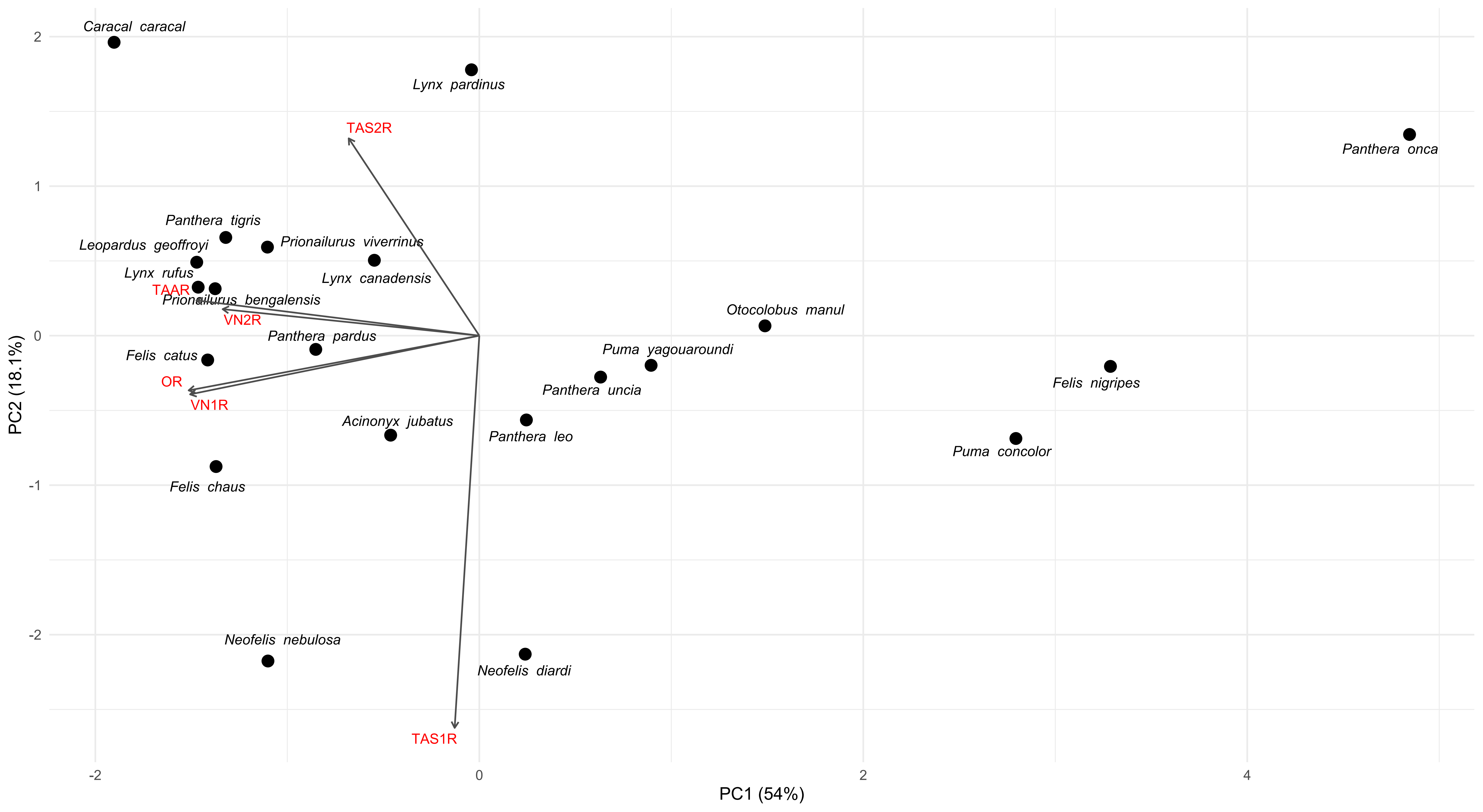

### Supplementary Figure 2

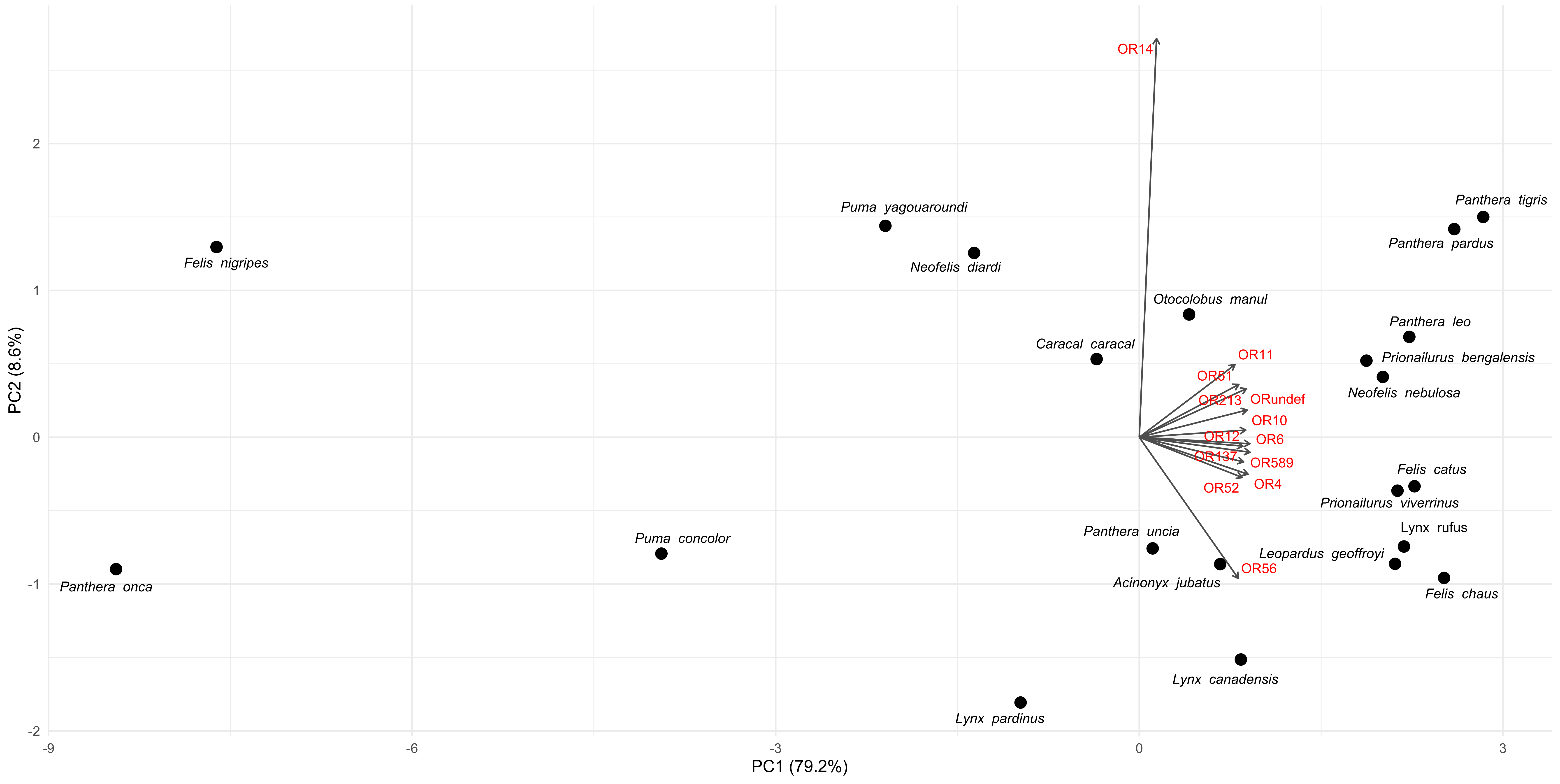

### Supplementary Figure 3

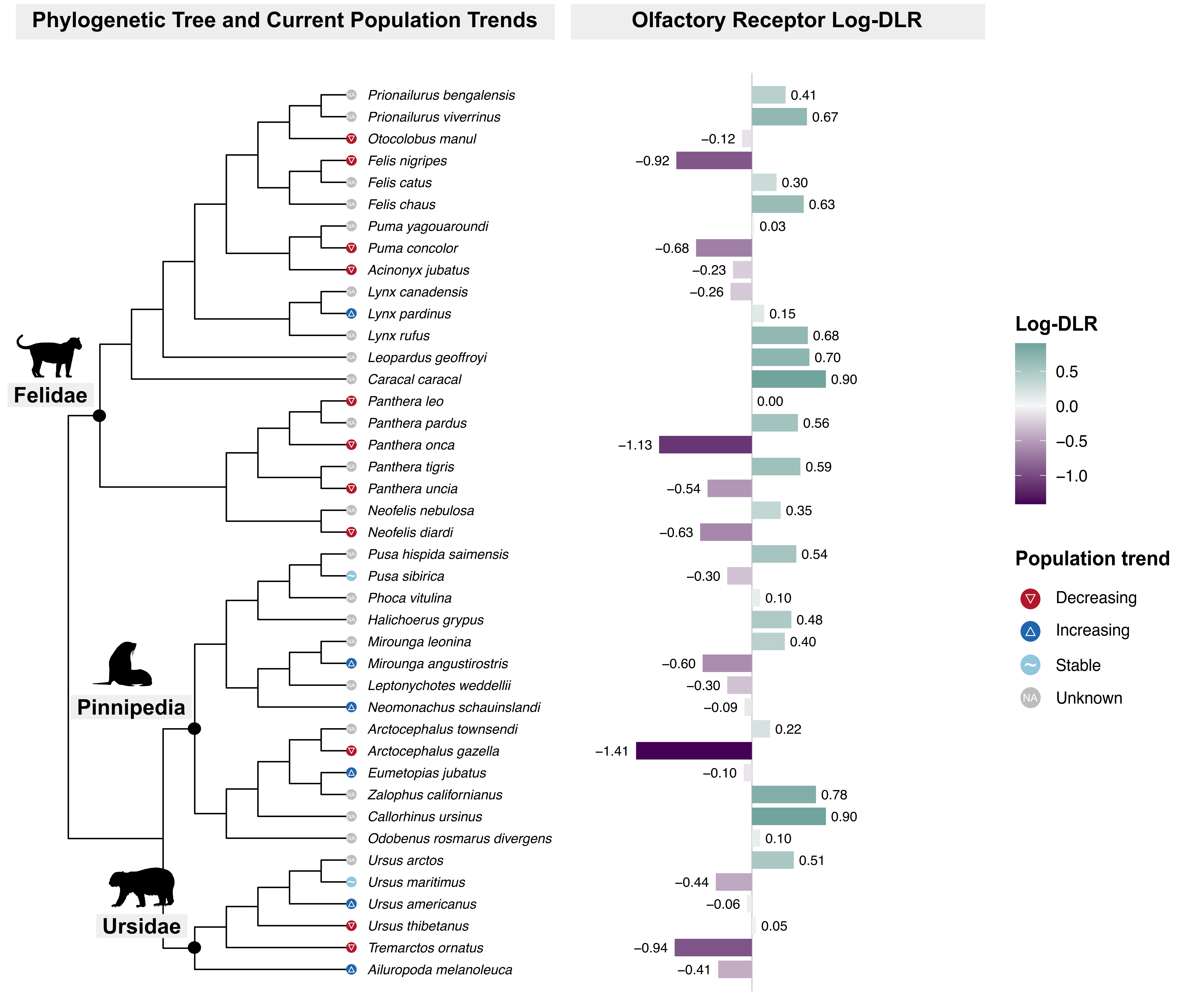
